# Optimal foraging models reveal barriers and stepping stones to the evolution of host-range generalism in bacteriophage

**DOI:** 10.1101/079889

**Authors:** Jeremy Draghi

**Affiliations:** Dept. of Biology, Brooklyn College; The Graduate Center, City University of New York

## Abstract

The nature and stability of coexistence of specialist species with more generalized competitors present theoretical questions that have been difficult to resolve. Recent surveys of bacteriophage host-ranges suggest that generalist phage often coexistent with specialists. However, previous theoretical work has explained this coexistence only in terms of strict genetic trade-offs, which are not consistently observed when phage are challenged to evolve to multiple hosts in laboratory environments. Here we use the framework of optimal foraging to identify conditions that might prevent generalists from outcompeting specialist relatives. Our analysis shows that heterogeneities in phage life-history properties make host-range specialist more viable, and that endogenous fluctuations in host density permit a narrow window of stable coexistence between specialists and generalists without the need for genetic trade-offs. These results are especially relevant for understanding the barriers to the evolution of broader host-ranges in bacteriophage and other pathogens with similar life-cycles.

## INTRODUCTION

The goal of understanding competition and coexistence among generalists and specialists has attracted significant theoretical effort in ecology (Wilson & Yoshimura 1994; Abrams 2006a). The complexity of this question has only grown as more recent work has highlighted the role of evolutionary dynamics in coexistence (Yoshida et al. 2003; Urban & Skelly 2006; Lankau 2011; Vasseur et al. 2011; Kremer & Klausmeier 2013) or debated the stability of generalist-specialist ecosystems over evolutionary time (Egas et al. 2004; Abrams 2006b). The resulting confusion is particularly acute in dynamic scenarios, such as invasions of introduced species or emergence of new pathogen strains, in which ecological coexistence with existing types can provide a foothold for adaptation and further expansion.

Bacteriophages and their hosts provide an ideal testbed for assessing and developing these diverse models of generalist-specialist coexistence. Phage host-ranges are both extremely diverse in extent (Weitz et al. 2013) and evolutionarily labile, showing large sensitivity to single mutations (Duffy et al. 2007) and evolving rapidly in the lab (Marston et al. 2012; Meyer et al. 2012). Evolution experiments have shown that host range mutants can displace (Bono et al. 2012) or coexist alongside their immediate ancestral phenotypes (Chao et al. 1977; Bono et al. 2015). On a larger scale, phage host-ranges may play a critical role in shaping bacterial diversity, and phage host-shifts can serve as a useful model for emergence and pathogenesis in human diseases.

Although phage have often been regarded as specialists, more recent surveys have revealed a complex pattern of nested diversity (Flores et al. 2011, 2013; Weitz et al. 2013), meaning that specialists coexist with generalists with overlapping host ranges. Ecological models have explored the stability and assembly of such communities and concludied that nested infection networks can arise if the fitness of phage with broader host range is restricted by genetic trade-offs—specifically, if fitness on a given host is negatively correlated with the breadth of host-range (Jover et al. 2013; Korytowski & Smith 2015). While such trade-offs are intuitive and sometimes observed (Duffy et al. 2006), evolution experiments have often produced generalists without compromised performance in comparison to specialists (Remold 2012; Bono et al. 2013). These observations might be artifacts of artificial environments or lab-adapted model systems but, when taken at face value, present a puzzle: if phage can readily evolve broader host ranges without apparent trade-offs, why do specialists persist?

Optimal foraging theory can provide a different perspective on phage host-ranges which may help resolve this puzzle. Such models typically focus on heterogeneity in the value of diverse hosts or other resources, rather than assuming direct costs to generalism. Because phage which infect a poor host necessarily lose any opportunity to instead infect a better one, such optimality models could predict picky host-ranges based on opportunity costs, even in the absence of genetic trade-offs in performance across hosts. Bull (2006) established a simple rule for host-range in lytic phage populations with no net growth: it is beneficial to infect any host whose effective burst size (i.e., devalued by the probability of host death before lysis) exceeds one. Intuitively, this rule arises because the reproductive value of a phage virion must be one if it has achieved steady-state on its original host (λ = 0) and if its environment is homogenous. While this result appears to set a low bar for the evolution of generalism, experiments have verified that phage can evolve to avoid nonproductive hosts (Heineman et al. 2008), and a recent model (Sieber & Gudelj 2014) shows that host defenses against phage (Labrie et al. 2010) may reduce effective burst sizes to values near one. In growing phage populations, these opportunity costs are more potent and phage can optimally evolve more discriminating host-ranges (Bull 2006; Guyader & Burch 2008). Here we model how violations of other key assumptions of Bull (2006) change optimal host-ranges in the absence of direct costs of generalism. Specifically, we focus on heterogeneity among phage virions and dynamics in persistently cycling populations, showing that each of these aspects of phage biology can significantly alter selection on the use of marginal alternative hosts.

## MODELS & RESULTS

### Scope & Terminology

We focus on bacteriophage, using “phage” to refer to a type or class and “virion” to refer to an individual infectious particle. These results should generalize to other viruses that diffuse through a liquid environment and can potentially encounter multiple possible hosts; this category includes viruses of single-celled eukaryotes in liquid environments as well as human pathogens encountering multiple cell-types as they move throughout the host.

### I. Heterogeneity among phage particles can select for a more narrow host-range

We begin with a model of a single phage type in an environment with two potential hosts, *A* and *B*. We assume that the phage initially only infects host *A* and that its population has reached a dynamic equilibrium on that host (*λ*_*phage*_ = 0). Let β_A_ equal the effective burst size on host *A*—that is, the mean number of virions produced per infection, accounting for any probability of premature cell death or resistance to the infection. The net yield of an infection on host A is then β_A_ minus one to represent the loss of the infecting particle itself. Let γ equal the rate of phage mortality when outside of a host, and let *θ*_*A*_ equal the per-capita rate of successful infection. *θ*_*A*_ can be decomposed into the expression *α*_*A*_ *N*_*A*_ / *V*, where *α*_*A*_ is the per-phage attachment rate (with units ml/(cells x minute)), *N*_*A*_, the abundance of the host cells, and *V*, the culture volume in milliliters.

Each virion must have an expected reproductive success, *W*, of one. The relation among these parameters is therefore:

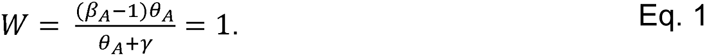

If we make the additional assumptions that phage particles do not senesce and the phage and hosts are well-mixed, and ignore complications like multi-step infection processes, then Eq. 1 holds for any phage particle outside of a cell. This expected fitness of one quantifies the opportunity that is forgone if a phage particle were to infect a cell of host *B*; that is, the opportunity cost of infecting *B*. Bull (2006) finds that, at equilibrium, infection of a host is favored if the yield of that host exceeds one; it is equivalent to note that the payoff of “investing” in host *B* must exceed the opportunity cost of *W* for the investment to be beneficial.

The value of using opportunity cost as a framework is seen when we deviate from the above assumptions. Consider a situation in which phage particles have non-heritable differences in their construction, such that some decay faster than others. How would this phenotypic variation affect the break-even value of an alternative host? Figure 1(a) frames the problem: a phage which has encountered a cell of host *B* could infect that cell, forgoing the chance of a later encounter with the superior host *A*. Because the high- and low-mortality types are not heritable, they cannot evolve separate strategies; the optimal strategy must be a compromise of the optima for each type. However, Fig. 1(b) illustrates why that compromise might be biased by heterogeneous life-history traits. Although, in this example, long-lived phage are equal in number to short-lived, their longer lives mean that each is more likely to encounter a cell of host *B*. Therefore, a phage encountering a cell of host *B* is often of the long-lived type, especially when host *B* is rare.

**Figure 1:**
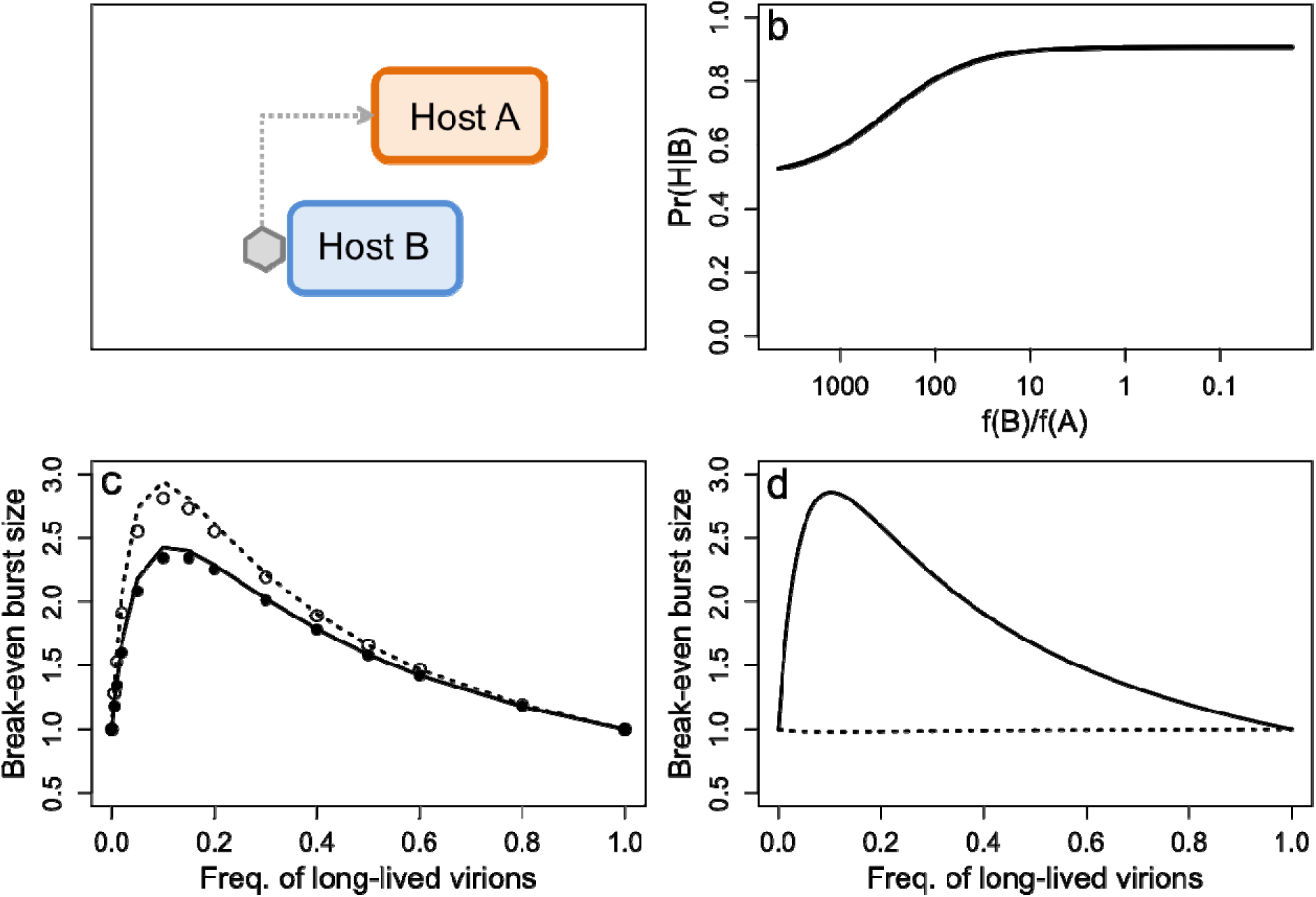
(a) Illustration of the predicament of a virion with a genetically determined host range. A virion that encounters a cell of host *B* could infect it, but would forgo a possibly of encountering the preferable host A. (b) The probability that a virion which encounters a cell of host *B* is of the long-lived type, given equal proportions of short- and long-lived types at lysis and the densities of host *B*, relative to the equilibrium density of host *A*, shown on the *x*-axis. (c) Effective burst size at which a broader host-range is neutral (break-even value) across a range of values for the frequency of phage particles with lower decay rates (type-*L*). *Solid, filled*: Carrying capacity of host *B* (*K*_*B*_) is 1 × 10^8^/ml; *dashed, open*: *K*_*B*_ is 1 × 10^7^/ml. Points represent results from numerical solutions to a differential equation model (see Appendix) while lines are solutions to Eq. 4. Parameters are as follows: *K*_*A*_ = 1 × 10^8^/ml, *β*_*A*_ = 100, *γ*_*L*_ = 0.001, *γ*_*H*_ = 0.01. (d) Effective burst size at which a broader host-range is neutral (break-even value) across a range of values for the frequency of phage particles with higher attachment rates. Lines are drawn from solutions to Eq. 5. *Solid*: attachment rate to host *B* is independent of the rate for host *A*; *dashed*: attachment rates to hosts *A* and *B* covary. Parameters are as follows: *K*_*A*_ = *K*_*B*_ = 1 × 10^8^/ml, *β*_*A*_ = 100, *γ* = 0.001, *θ*_*H*_ = 10θ_L_.

Let *p* represent the fraction of phage particles which have the lower decay rate, *γ*_*L*_, with 1-*p* having the worse decay rate *γ*_*H*_. Eq. 1 can then be adjusted to track the contributions of two classes of phage:

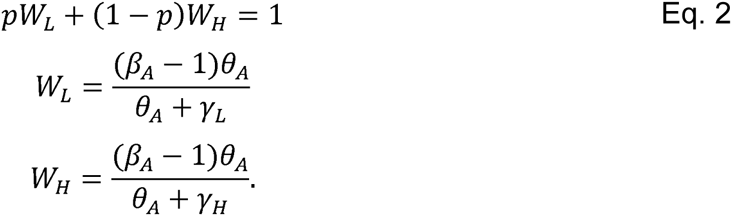

The *L* and *H* types differ in their opportunity costs; in particular, type-*L* should optimally infect host *B* only if its burst size on that host exceeds *W*_*L*_, which itself is greater than one if *γ*_*L*_ < *γ*_*H*_. Absent any mechanism by which *L* and *H* types could have distinct infection behaviors, the break-even benefit, *β\**, of host *B* must be an average of *W*_*L*_ and *W*_*H*_, weighted by the initial prevalence and longevity of each type. We can write this weighted average using expressions of the form *Pr(X | B)*—the probability that a particle is of type *X*, given that it encountered a cell of host *B*.

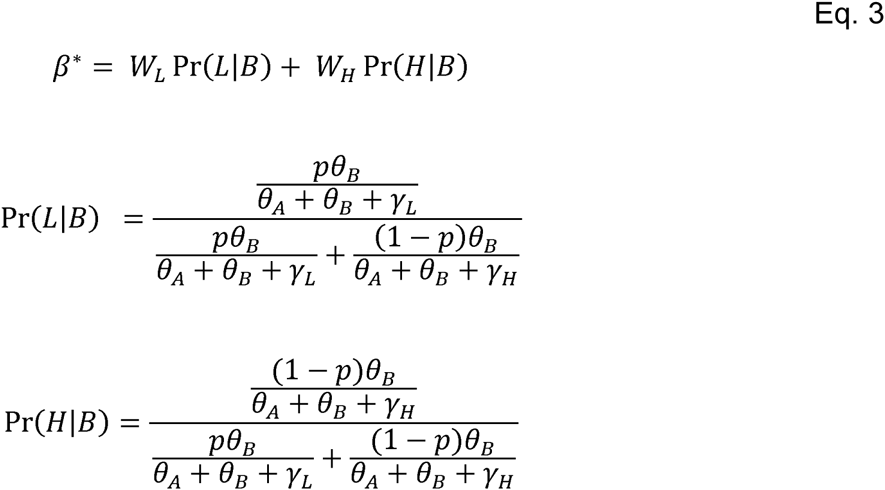

We can simplify this by noting that, if burst sizes are large, then at equilibrium most phage decay before finding a host. Therefore, if *β*_*A*_ >> 1, then *θ*_*A*_ << *γ*. Making the approximations *θ*_*A*_ + *γ*_*L*_ ≈ *γ*_*L*_, *θ*_*A*_ + *γ*_*H*_ ≈ *γ*_*H*_ and *β*_*A*_ −1 ≈ *β*_*A*_, We can more easily solve for *α*_*A*_ in Eq. 2 and obtain an approximate solution:

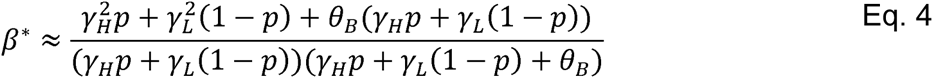

The predictions from this equation plotted in Figure 1(c) illustrate three main points. First, heterogeneity in death rates increases the break-even host values, skewing them substantially from the prior prediction of one. Second, in contrast to the simpler model explored in Bull (2006) and Heineman, Springman and Bull (2008), the density of a potential host can qualitatively alter whether it is beneficial to exploit that host. Intuitively, when the prevalence of a potential host is low, then the long-lived phage particles with high reproductive values compose a higher fraction of the particles that encounter that potential host (Fig, 1(b)). Finally, the approximation in Eq. 4 performs adequately when compared to numerical simulation of a differential-equation model of the system (Appendix A), validating the use of opportunity costs to predict these outcomes.

We also investigated the effects of nonheritable variation in attachment rates on optimal host-range. We derive an expression analogous to Eq. 4, this time retaining all *θ* terms to fully capture the effects of heterogeneity in this parameter. Let 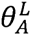 quantify the attachment rate of the low-attachment type and 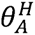 the rate for the high-attachment type. We can then write an equilibrium equation similar to Eq. 1, find equilibrium values of the attachment rates, and calculate *β\** as above.

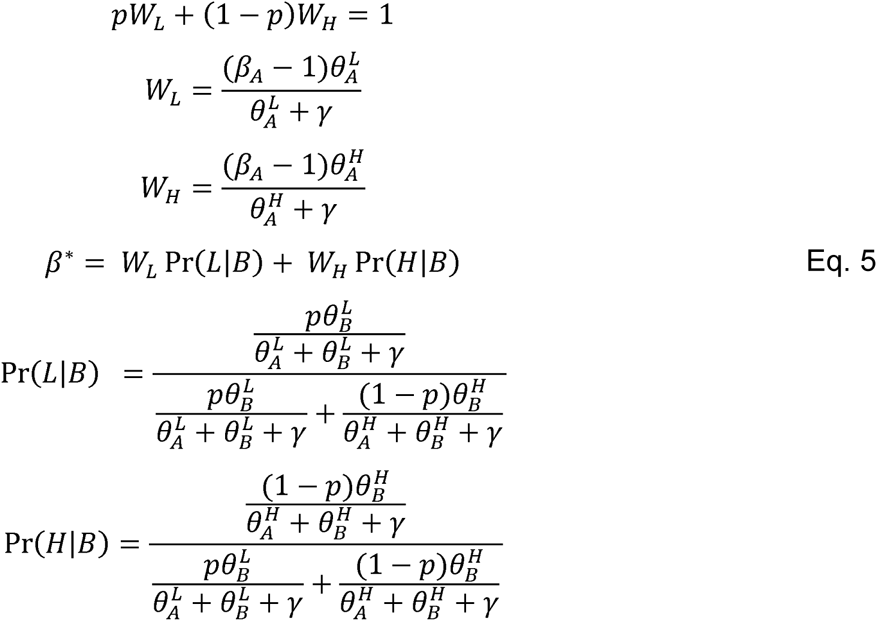

We can consider two models: in the first, phage attachment rate for host *A* varies while attachment for host *B* remains constant 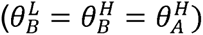, while in the second, 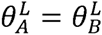 and 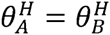. Figure 1(d) illustrates the substantial impact of this choice. When attachment rates across hosts covary, then the high-fitness, high-attachment-rate particles also have a proportionally higher chance of infecting host *B* cells. Therefore, the value of the alternative host must exceed a higher threshold to offset the lost reproductive potential of this high-attachment-rate particles.

### II. Phage latent period interacts with endogenous population cycles to allow host-range diversity

The results in Part I demonstrate that optimal host range can vary in equilibrium populations. However, phage populations may exhibit long-term persistence without equilibrium through stable oscillations or limit cycles (Levin et al. 1977; Weitz 2016). In order to capture a dynamic balance among phage and their hosts, we used a delay differential equation model. This framework allows an explicit delay called the latent period, *φ*, between infection and lysis. We model a well-mixed population where infections are the product of an attachment rate, *α*_*xy*_, the abundance *P*_*x*_ of phage *x* and the abundance *N*_*y*_ of host *y*. For simplicity, we model two phage types: a specialist *P*_*S*_ which only infects host *A*, and a generalist *P*_*G*_ which infects both host *A* and *B* with equal attachment rates. To focus on host-range generalism without explicit costs, we stipulate that *α*_*SA*_ = *α*_*GA*_ = *α*_*GB*_ and therefore omit subscripts on *α*. Phage decay at a rate *γ* while hosts grow according to Lotka-Volterra dynamics with parameters *r*_*x*_ and *K*_*x*_ for host *x*. We also model three classes of infected hosts, corresponding to each of the combinations of phage and host. The model is similar to Weitz (2016, p. 82), though without an explicit treatment of host mortality.

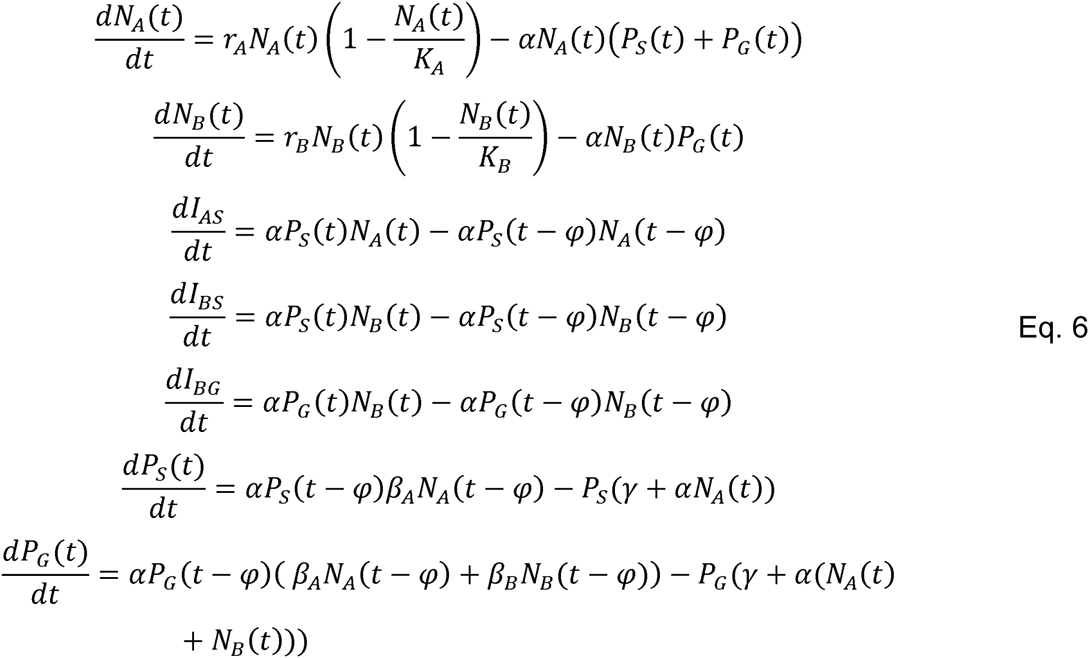

Figure 2(a-c) depicts the long-term behavior of systems with only the specialist phage and host *A*. Significant latent periods induce periodic dynamics, with a longer latent period leading to higher amplitude cycles. Figure 2(d) shows the effects of this parameter change on the success of host-range generalists. When *φ* = 0, the results conform perfectly to Bull (2006): generalists are favored when the effective burst size on host *B* is greater than one and disfavored otherwise. However, at *φ* = 30 and *φ* = 100, the outcomes change qualitatively. Generalist phage exhibit ecologically stable coexistence with specialists for values of host *B* near one.

**Figure 2:**
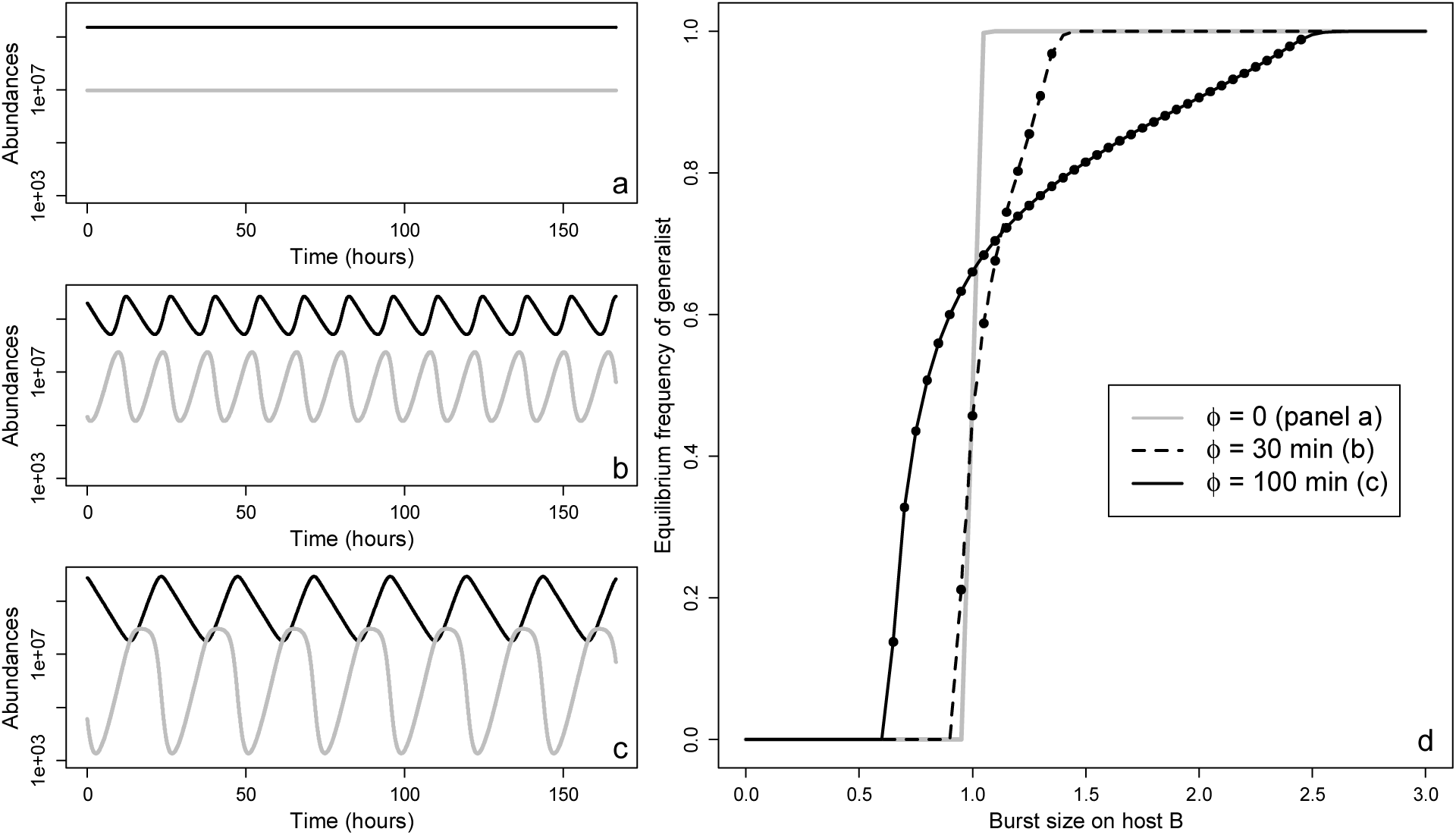
(*a-c*) Long-term behavior of the abundances of specialist phage (*black*) and host (*grey*) when the latent period *φ* equals zero (*a*), 30 minutes (*b*), and 100 minutes (*c*). Other parameters are: *r* = 0.02, *K* = 1 × 10^8^, *m* = 0.0075, β_A_ = 100, and α = 8 × 10^−12^ min^−1^. (d) Mean long-term frequency of generalists across values of β_B_ for *φ* = 0, 30, and 100 minutes. Means are averaged over a window of 10,000 minutes at the end of one million minutes of forward-time simulation; for each point, a simulation with an initial frequency of 1 × 10^−6^ generalists is averaged with one in which specialists, rather than generalists, are initially rare. Filled circles depict tested values for which both generalists and specialists increased when rare; other points are elided.

To further explore this result, we examined the fine-grain dynamics of competition between specialist and generalist phage over a single cycle. The phage populations grow when their host are at high density; during this phase of a population cycle, host-range specialists have an advantage (Fig. 3). Exhaustion of host *A* and, to a lesser degree, host *B*, then precipitates a net decline in the phage population. During this phase, host-range generalists have an advantage, lasting until the population of host *A* has nearly recovered.

**Figure 3:**
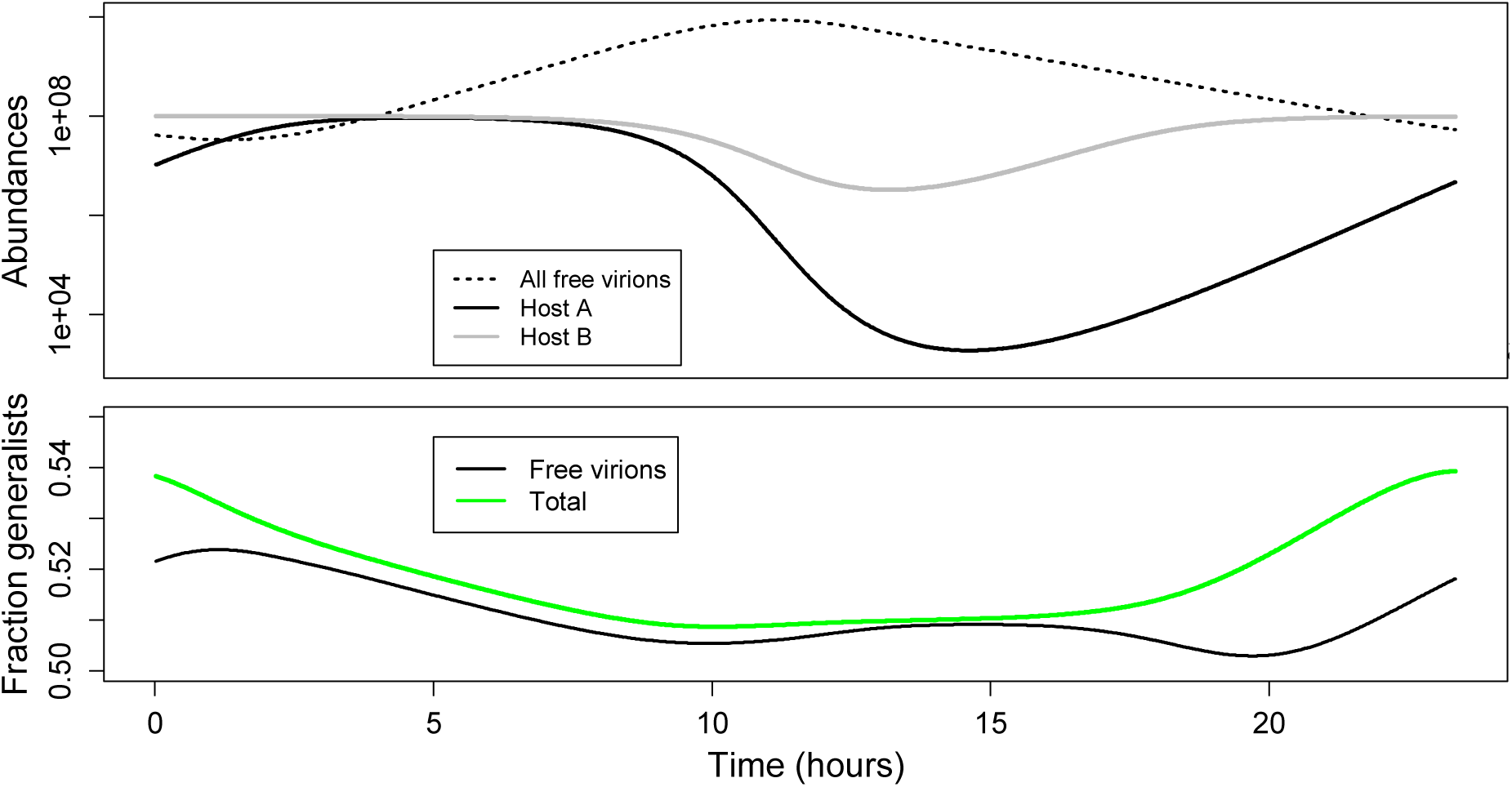
Detailed dynamics over one cycle of coexisting host-range generalists and specialists at their dynamic equilibrium. The green line represents the ratio of generalists to all phage when the fitness contributions of current infections are totaled together with free virions. Note that this measure of relative success yields a simple pattern: generalists do better than specialists when the population of free virions is falling and worse when the number of free virions is growing. Parameters are *φ* = 100 min, *r* = 0.02, *K* = 1 × 10^8^, *m* = 0.0075, β_A_ = 100, [check beta B] and α = 8 × 10^-12^ min^-1^.

While Fig. 3 shows an equilibrium between specialists and generalists, the mechanism of stable coexistence is not apparent. We hypothesized that the impact of generalists on the density of host *B* limits their success: if generalists become more common, then host *B* would crash more severely, reducing the benefit of generalism without affecting its cost. Mathematically, we can test this hypothesis by removing the phage mortality term, *αN*_*B*_(*t*)*P*_*G*_(*t*), from Eq. 6. With mortality, generalists increase when common (i.e., above 99% of the population) only when β_B_ exceeds approximately 2.5; without mortality, this threshold drops to approximately 0.65.

Large values of the latent period have two significant effects in the simulations shown in Fig. 2: they induce endogenous fluctuations but also effectively shelter an infecting virion from decay for the duration of the latent period. Figure 3 illustrates the benefit of the latter effect that accrues to generalists during the period of net negative growth. However, the results in Part I predict that non-heritable heterogeneity in attachment and decay rates will also affect selection on generalists, suggesting that fluctuations per se might also help or hinder generalists. To test this, we simulated exogenous fluctuations in host growth rates in a parameter regime without endogenous fluctuations. Each growth rate in Eq. 6 was replaced with the expression:

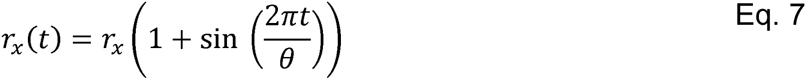

where θ is the period in minutes.

Figure 4 shows that exogenous fluctuations can drive generalist-specialist coexistence, but only when *φ* > 0. This result confirms the importance of modeling a finite, rather than implicit, latent period and generalizes the type of fluctuations that can skew the optimal host-range from the expected outcome.

**Figure 4:**
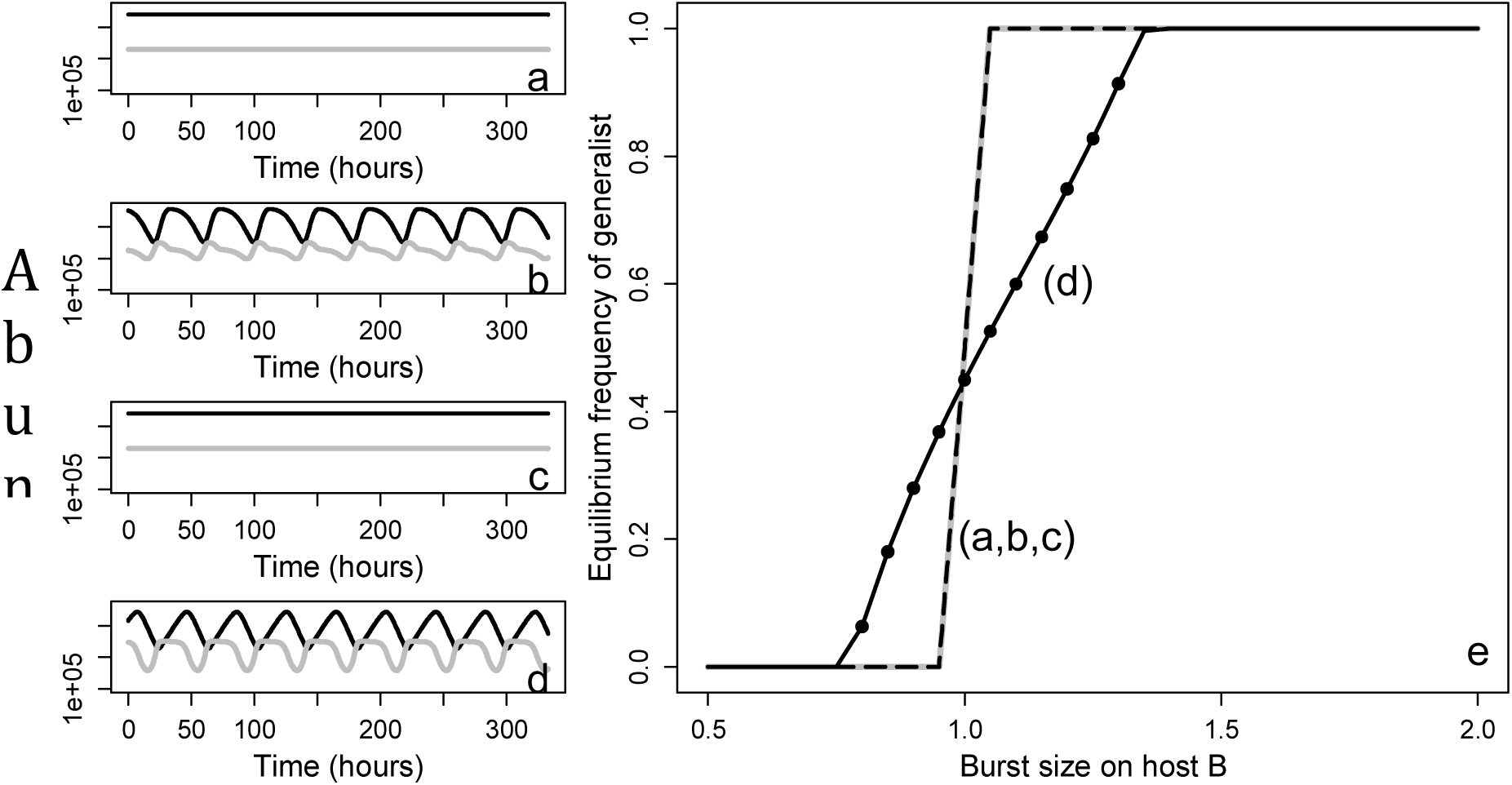
(*a-d*) Long-term behavior of the abundances of specialist phage (*black*) and host (*grey*) when the latent period *φ* equals zero (*a, b*) or 100 minutes (*c, d*) and exogenous change in host growths rates is absent (*a,c*) or present (*b,d*). Other parameters are: *θ* = 60 min*, r* = 0.02, *K* = 1 × 10^8^, *m* = 0.0075, β_A_ = 100, and α = 2 × 10^-12^ min^-1^; note that endogenous cycles are absent at this attachment rate. (e) Mean long-term frequency of generalists across values of β_B_. Note that lines for the systems illustrated in panels *a-c* overlap. Means are averaged over a window of 20,000 minutes at the end of five million minutes of forward-time simulation; for each point, a simulation with an initial frequency of 1 × 10^-6^ generalists is averaged with one in which specialists, rather than generalists, are initially rare. Filled circles depict tested values for which both generalists and specialists increased when rare; other points are omitted.

The existence of a stable mix of generalists and specialists suggests that a more flexible type of generalist, with independent rates of attachment to host *A* and *B*, might possibly invade to fixation. To test this hypothesis, we simulated a generalist, in competition with a specialist, with reduced rates of attachment to host *B* for several values of β_B_. This model is equivalent to Eq. 6 with the addition of a coefficient modifying the rate of attachment to *B*. In each tested case, a generalist with reduced affinity to the worse host can fully invade (Fig. 5).

**Figure 5:**
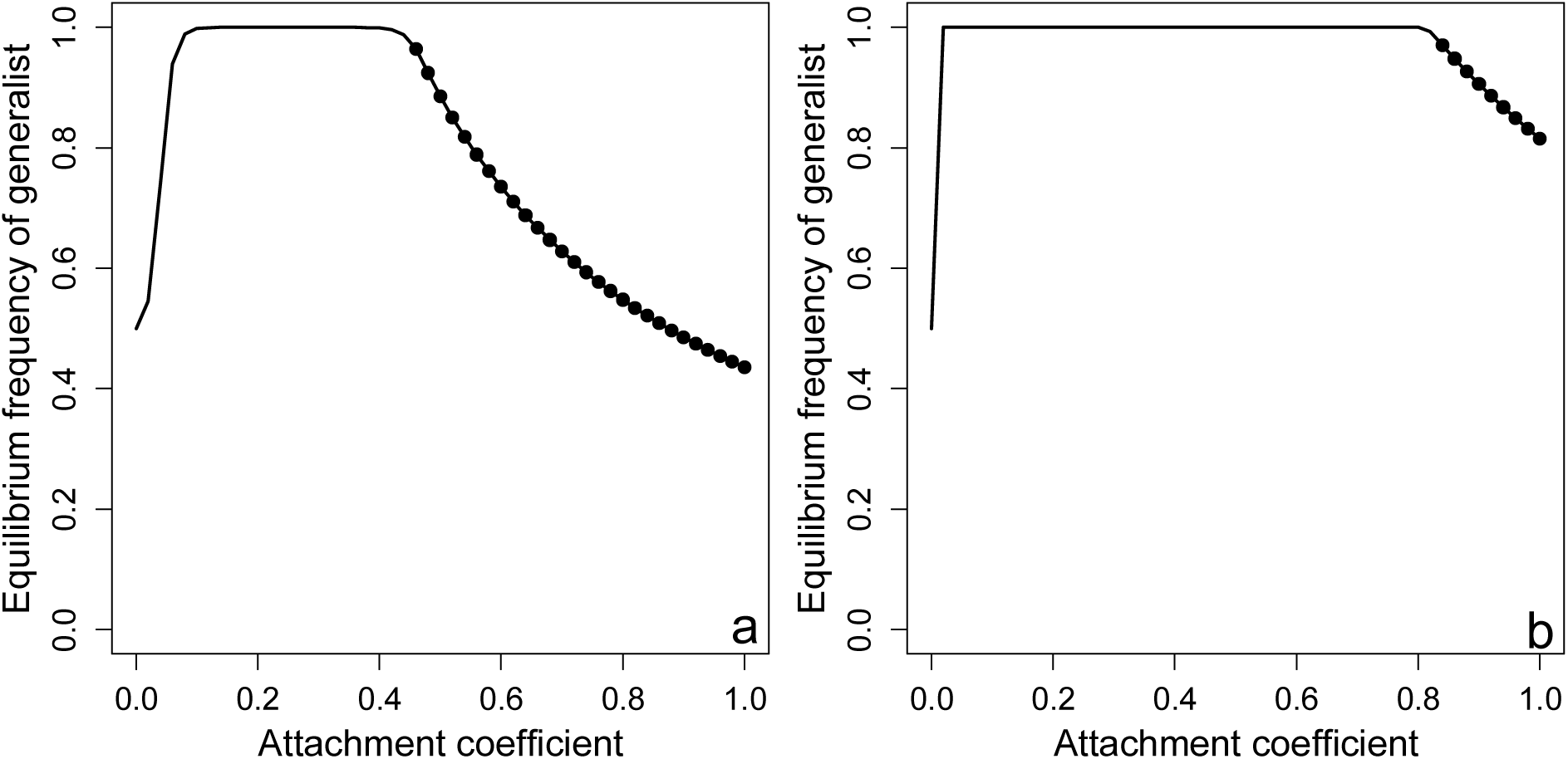
Mean long-term frequency of generalists across values the attachment rate to host *B*, relative to host *A*. Parameters are: *φ* = 100 min*, r* = 0.02, *K* = 1 × 10^8^, *m* = 0.0075, β_A_ = 100, and α = 8 × 10^-12^ min^-1^. Means are averaged over a window of 20,000 minutes at the end of one million minutes of forward-time simulation; for each point, a simulation with an initial frequency of 1 × 10^-6^ generalists is averaged with one in which specialists, rather than generalists, are initially rare. Filled circles depict tested values for which both generalists and specialists increased when rare; other points are omitted.

## DISCUSSION

The host ranges of viral pathogens of humans are notably dynamic. Because their hosts are capable of such rapid growth and evolution, bacteriophages must face an equally dynamic landscape of host choices. We propose that a better understanding of selection on the use of marginal hosts—hosts who, by virtue of poor resources, high defenses, or mismatched genetics, offer little return to infecting phage—is an essential step in predicting the evolution of broader host-ranges. Our results show that Bull (2006) is largely robust—in phage populations with sustainable dynamics, alternative hosts can be used optimally if their burst sizes exceed relatively low thresholds. While expanding this result well outside its original model, our results have also highlighted factors which skew these burst-size thresholds. In the process, we have found a type of coexistence among specialists and generalists which is stable even in the absence of direct costs of generalism.

Our models highlight the importance of explicitly considering the latent period in models of phage-host dynamics. Without a latent period, the opportunity costs that drive the generalist-specialist coexistence in part II disappear entirely. Our results depart significantly from those in Jover et al. (2013) due to this difference, as well as other differences, such as differing values of hosts, that reflect our focus on optimal use of marginal hosts. Without contradicting the results in Jover et al., our findings illustrate the need to match the diversity of phage life-histories with diverse approaches to modeling.

Experimental evidence for non-heritable heterogeneity in phage is relatively limited. Common phage model systems seem well-described by a homogenous process of exponential decay in lab enviroments (De Paepe & Taddei 2006). Heterogeneity in attachment rates has long been observed across many phage (Storms & Sauvageau 2014); this heterogeneity is non-heritable in lambda phage (Gallet et al. 2012) but not in T4 (Storms & Sauvageau 2014). Phage may also incorporate multiple genes expressing distinct and substitutable versions of a protein (Scholl et al. 2001), potentially leading to heterogeneity in mature, assembled virions. An equivalent type of heterogeneity could also arise in segmented phage with a high degree of co-infection; a polymorphism on one segment could create phenotypic variation from the perspective of unlinked loci on other segments. Heterogeneity in phage may be an unavoidable result of defects in protein folding or capsid assembly or an adaptive, bet-hedging response to environmental uncertainty (Gallet et al. 2012). Our results provide an additional impetus to characterize this form of phenotypic variation as a factor in predicting the optimality of life-histories and the probability of host-range shifts.

While this work provides some additional evidence of theoretical barriers to generalism, it is difficult to see how optimal foraging alone can explain the existence of specialists in systems in which generalists suffer few direct costs. Other promising ideas, such as epistatic barriers to the evolution of broader host-ranges (Remold 2012) or evolutionary costs of generalism (Whitlock 1996; Bono et al. 2015), should be integrated with optimal foraging approaches in future work.

## APPENDIX Model for Heterogeneous Decay Rates

Parameters are as defined above, except that both specialist and generalist phage are divided into high-mortality (P_SH_ and P_GH_) and low-mortality (P_SL_ and P_GL_) types.

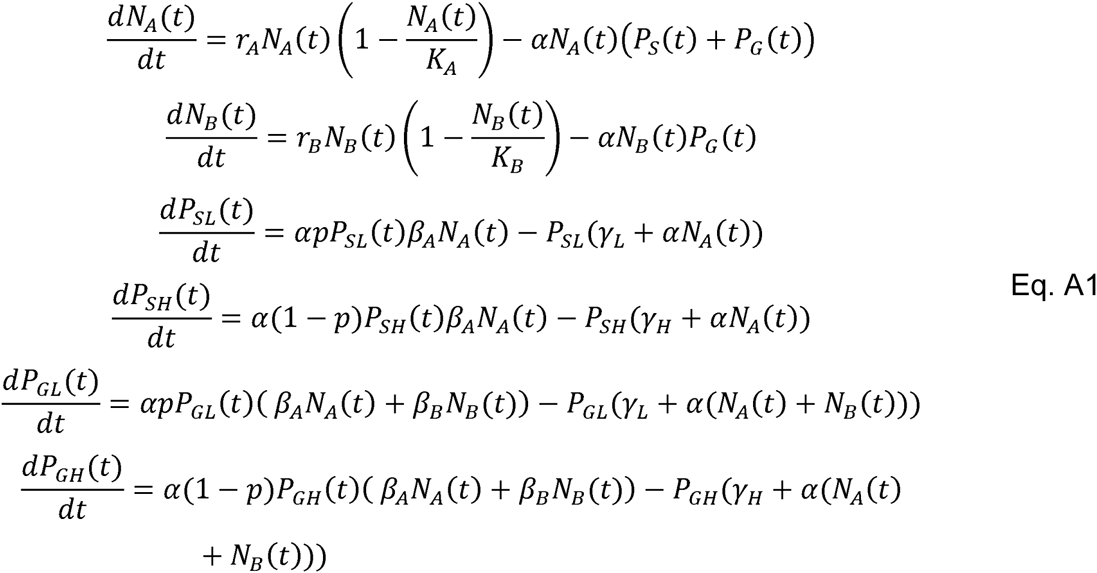

